# Studies in alkaptonuria reveal new roles beyond drug clearance for phase I and II biotransformations in tyrosine metabolism

**DOI:** 10.1101/2020.04.16.044347

**Authors:** Brendan P Norman, Andrew S Davison, Juliette H Hughes, Hazel Sutherland, Peter J Wilson, Neil G Berry, Andrew T Hughes, Anna M Milan, Jonathan C Jarvis, Norman B Roberts, Lakshminarayan R Ranganath, George Bou-Gharios, James A Gallagher

## Abstract

**Background and Purpose:** alkaptonuria (AKU) is an inherited disorder of tyrosine metabolism caused by lack of the enzyme homogentisate 1,2-dioxygenase (HGD). The primary biochemical consequence of HGD-deficiency is increased circulating homogentisic acid (HGA), which is central to AKU disease pathology. The aim of this study was to investigate the wider metabolic consequences of targeted *Hgd* disruption.

**Experimental Approach:** the first metabolomic analysis of the *Hgd*^−/−^ AKU mouse model was performed. Urinary metabolites altered in *Hgd*^−/−^ were further validated by showing that the HGA-lowering drug nitisinone reversed their direction of alteration in AKU

**Key Results:** comparison of *Hgd*^−/−^ (AKU) versus *Hgd*^+/−^ (heterozygous control) urine revealed increases in HGA and a group of 8 previously unreported HGA-derived transformation products from phase I and II metabolism. HGA biotransformation products HGA-sulfate, HGA-glucuronide, HGA-hydrate and hydroxymethyl-HGA were also decreased in urine from both mice and patients with AKU on the HGA-lowering agent nitisinone. *Hgd* knockout also revealed a host of previously unrecognised associations between tyrosine, purine and TCA cycle metabolic pathways.

**Conclusion and Implications:** AKU is rare, but our findings further what is currently understood about tyrosine metabolism more generally, and show for the first time that phase I and II detoxification is recruited to prevent accumulation of endogenously-produced metabolites in inborn errors of metabolism. The data highlight the misconception that phase I and II metabolic biotransformations are reserved solely for drug clearance; these are ancient mechanisms, which represent new potential treatment targets in inherited metabolic diseases.

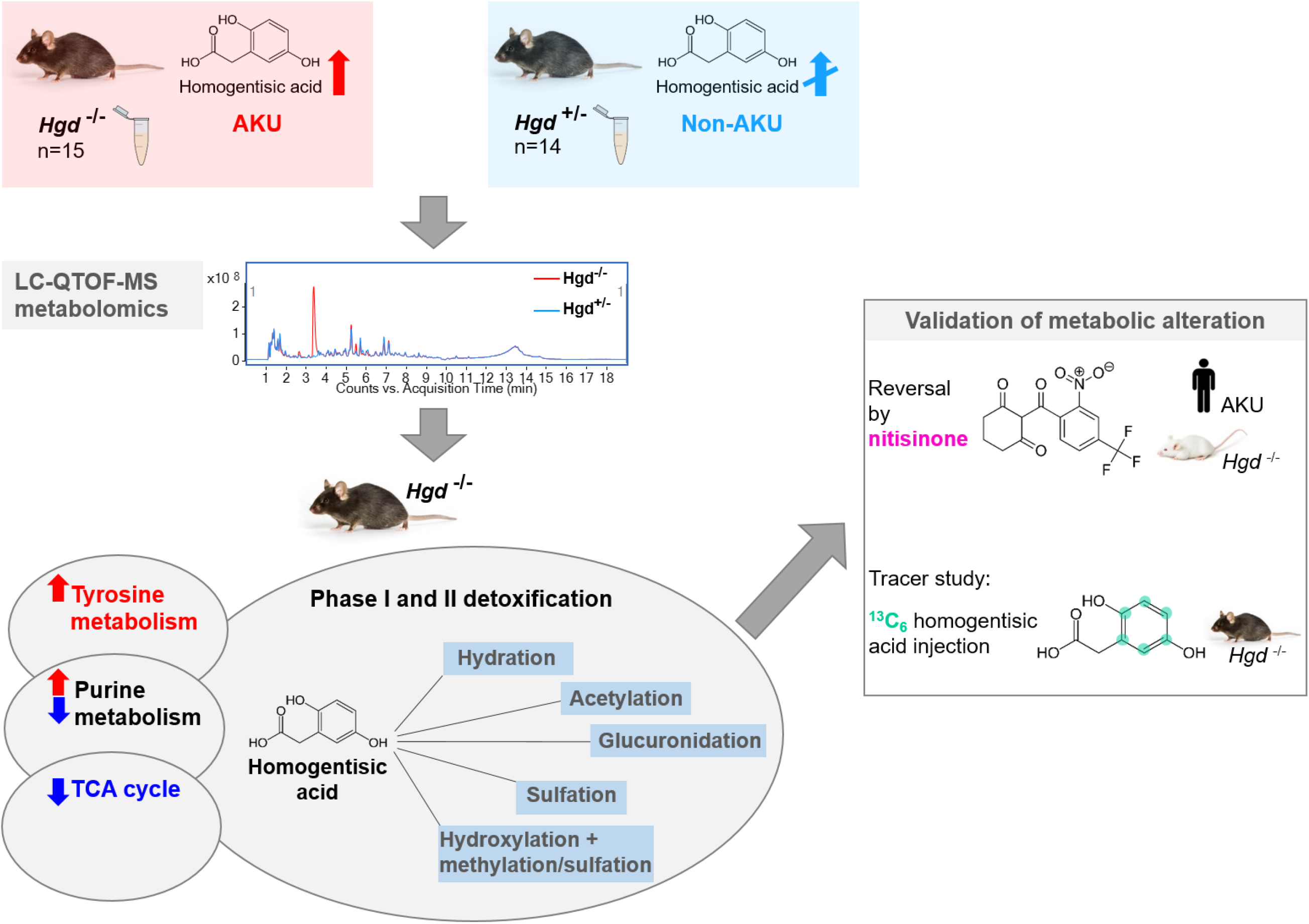

**Bullet point summary:** What is already known

- Increased circulating homogentisic acid is central to disease pathology in the inherited metabolic disease alkaptonuria
- The *Hgd* knockout mouse, created in our laboratory, accurately models human alkaptonuria

What this study adds

- Phase I and II biotransformations are recruited in alkaptonuria for detoxification of homogentisic acid
- These data challenge misconceptions that phase I and II metabolism is solely for drug clearance

Clinical significance

- Phase I and II metabolic processes represent new treatment targets in inherited metabolic diseases
- The molecular pathology of AKU extends much further than the known alteration to tyrosine metabolism

## 1. Introduction

Alkaptonuria (AKU) is a rare disorder of tyrosine metabolism caused by congenital lack of the enzyme homogentisate 1,2-dioxygenase HGD (E.C.1.12.11.5) (Zatkova, 2011). The biochemical consequence of this deficiency is increased homogentisic acid (HGA) in the circulation, the pathognomonic sign of the disease which is central to its pathophysiological features (Ranganath et al., 2013). HGA has a high affinity for collagenous tissues, where its deposition produces striking pigmentation (Helliwell et al., 2008), a process called ochronosis. Cartilage of load-bearing joints is particularly susceptible to ochronosis. Presence of HGA-derived pigment in these joints alters the physicochemical properties of cartilage that support normal transmission of load and results in an inevitable and severe early-onset osteoarthropathy (Taylor et al., 2011; Chow et al., 2020).

Metabolomics has emerged as an invaluable approach for studying AKU. Mass spectrometric assays have been developed as a targeted metabolomic approach for diagnosis and monitoring of AKU (Hughes et al., 2014, 2015a). These assays offer precise quantitation of tyrosine pathway metabolites including HGA in serum and urine. More recently, a strategy for profiling and chemical identification of hundreds of metabolites simultaneously by liquid chromatography quadrupole time-of-flight mass spectrometry (LC-QTOF-MS) has been developed (Norman et al., 2019). Application of this technique to AKU serum (Davison et al., 2019a) and urine (Norman et al., 2019) enabled the discovery of previously unknown metabolite and metabolic pathway alterations following treatment with the HGA-lowering agent nitisinone.

Metabolic profiling has potential in AKU as both a phenotyping and biomarker discovery tool. However, to our knowledge, untreated AKU has not been compared with non-AKU at the metabolome level before. Here we compared the metabolome of AKU mice, a model created by targeted homozygous knockout of the *Hgd* gene (*Hgd*^−/−^), with non-AKU heterozygous knockout (*Hgd*^+/−^) control mice (Experiment 1). The *Hgd*^−/−^ mouse recapitulates human AKU, with elevated plasma and urine HGA, development of ochronosis and its inhibition by nitisinone (Preston et al., 2014; Hughes et al., 2019). We then looked to validate any metabolite alterations observed by assessing whether the direction of alteration was reversed while on the HGA-lowering drug nitisinone treatment in *Hgd*^−/−^ mice or patients (Experiment 2), and whether the metabolites that were increased in *Hgd*^−/−^ mice were directly derived from HGA via a metabolic flux experiment in which mice were injected with stable isotopically-labelled HGA (Experiment 3). Figure 1 summarizes the overall study design. The overall aim of these experiments was to investigate metabolic markers of disease in AKU; identification of novel metabolite alterations in a controlled mouse study could elucidate novel disease mechanisms and improve current strategies for monitoring the disease process and treatment response.

**Figure 1.**
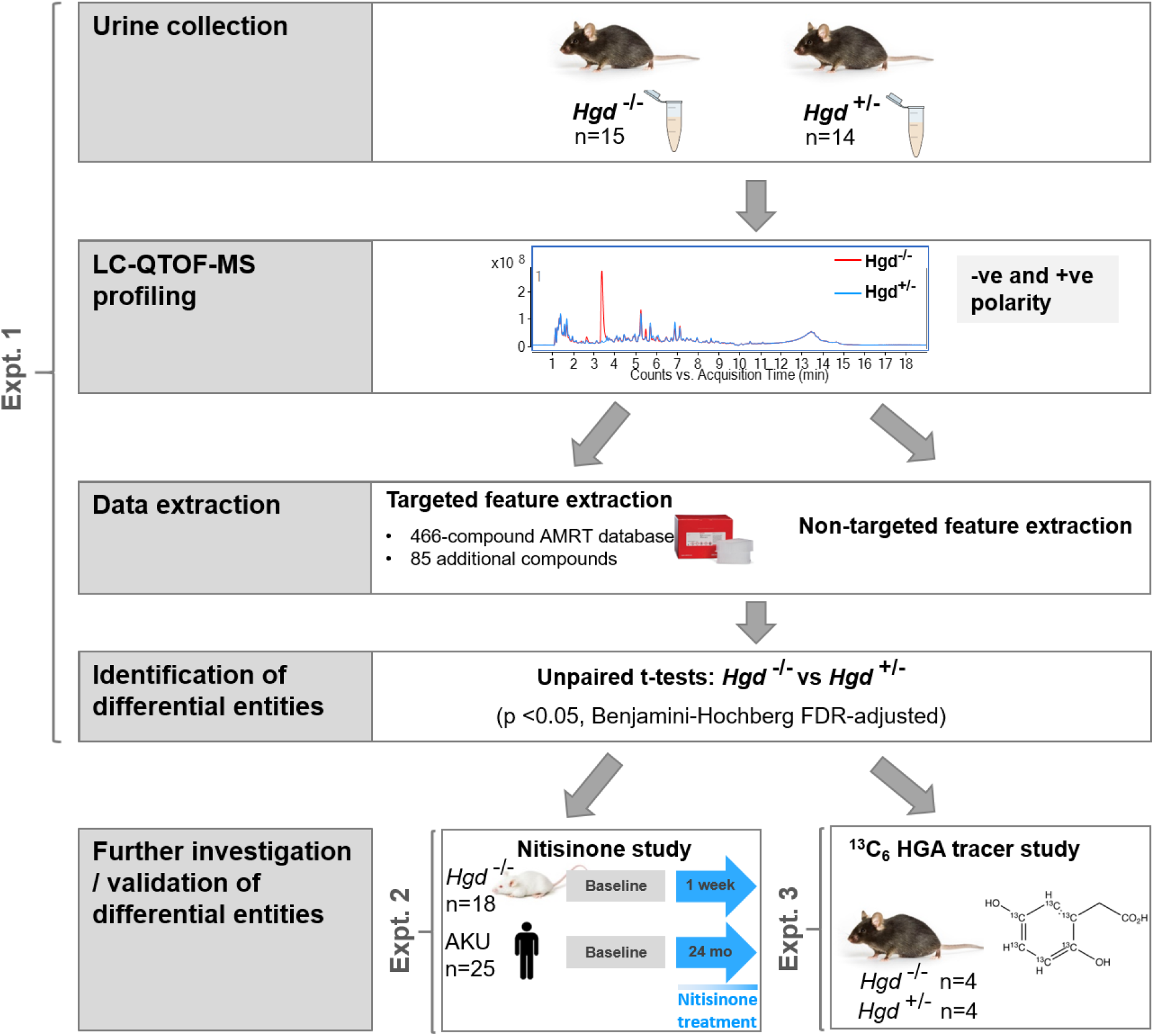
Schematic overview of the overall study design, incorporating Experiments 1-3. In Experiment 1, urine was collected from *Hgd*^−/−^ and *Hgd*^+/−^ mice and profiled by LC-QTOF-MS. Targeted and non-targeted feature extraction was performed on the data in parallel and subsequent unpaired t-tests were employed to identify differentially abundant chemical entities between *Hgd*^−/−^ and *Hgd*^+/−^. These entities were then further investigated in LC-QTOF-MS data from two additional datasets; a previously published study examining the effect of nitisinone on the urine metabolome of *Hgd*^−/−^ BALB/c mice and patients with AKU (Norman et al., 2019) (Experiment 2) and a plasma flux analysis using a ^13^C_6_ labelled HGA tracer (Experiment 3).

## 2. Methods

### 2.1 Experimental Design

The study comprised three experimental components (Figure 1). In Experiment 1 we employed non-targeted LC-QTOF-MS profiling to characterize the metabolome of a mouse model of AKU by targeted homozygous *Hgd* knockout (*Hgd*^−/−^), with non-AKU heterozygous knockout (*Hgd*^+/−^) mice as controls (Experiment 1). We then looked to validate the metabolite alterations observed by assessing whether the direction of alteration was reversed following nitisinone treatment in *Hgd*^−/−^ mice or patients with AKU (Experiment 2), and additionally whether the metabolites that were increased in *Hgd*^−/−^ mice were directly derived from the increased HGA via a metabolic flux experiment in which mice were injected with stable isotopically-labelled HGA (Experiment 3). All mice were housed in the University of Liverpool’s Biomedical Services Unit under pathogen-free conditions, in cages of up to 5 mice, with 12-hr light/dark cycle, and food and water available *ad libitum*. Mice were drug/test naïve at baseline in each experiment.

### 2.2 Materials

Deionised water was purified in-house by DIRECT-Q 3UV water purification system (Millipore, Watford, UK). Methanol, acetonitrile, isopropanol (Sigma-Aldrich, Poole, UK), formic acid (Biosolve, Valkenswaard, Netherlands) and ammonium formate (Fisher Scientific, Schwerte, Germany) were LC/MS grade. ^13^C_6_ labelled HGA for metabolic flux analysis was purchased from Toronto Research Chemicals (Toronto, Canada).

### 2.3 Urine sample collection and preparation for metabolic profiling (Experiment 1)

For metabolomic analysis of the targeted *Hgd* knockout phenotype (Hughes et al., 2019), urine was collected from 15 *Hgd*^−/−^ (mean age ±SD 12.8±0.1 weeks) and 14 *Hgd*^+/−^ (mean age 11.6±0.3) weeks) male C57BL/6 mice. All urine was taken on a single-collection basis and stored at −80°C.

Pooled samples were created in each profiling experiment for quality control (QC) purposes. For *Hgd*^−/−^ and *Hgd*^+/−^ groups a separate representative pool was created by pooling 20μL of each individual urine sample. An overall pool was also created for each experiment by pooling equal proportions of the above group pools. Analysis of individual and pooled samples was performed following dilution of 1:9 urine:deionised water as previously described (Norman et al., 2019).

### 2.4 Investigating the effect of nitisinone on metabolites showing alteration in *Hgd*^−/−^ mice and patients with AKU (Experiment 2)

The effect of nitisinone treatment on urinary metabolites altered in *Hgd*^−/−^ mice (Figure 1, Experiment 1) was studied in the data from a previous profiling experiment described by Norman *et al*. (Norman et al., 2019). These data were from urine from 18 BALB/c *Hgd*^−/−^ mice (mean age 27±12 weeks, 9 female, 9 male) with *Hgd* disruption by ENU mutagenesis (Preston et al., 2014) and from 25 patients attending the UK National Alkaptonuria Centre (NAC; mean age 51±15 years, 13 female, 12 male). The disease phenotypes of *Hgd*^−/−^ mice from targeted knockout and ENU mutagenesis models are identical (Hughes et al., 2019). Mouse urine was collected at baseline then after 1 week on nitisinone, administered in drinking water at 4mg/L, supplied *ad libitum*. Human urine was collected at baseline then after 24 months on nitisinone (2mg daily). These datasets were acquired under identical LC-QTOF-MS analytical conditions to those employed in the present study.

### 2.5 Design of in vivo metabolic flux experiment and sample collection (Experiment 3)

Eight C57BL/6 mice were studied in the HGA metabolic flux experiment; 4 *Hgd*^−/−^ (mean age 56±2.3 weeks, 1 female, 3 male) and 4 *Hgd*^+/−^ (mean age 58±0 weeks, 4 female). A 1.96mg/mL ^13^C_6_ HGA tracer solution was prepared in sterile saline and injected into the tail vein. Injection volume was adjusted for each mouse to achieve a final blood concentration of 1mmol/L, assuming a total blood volume of 75mL/kg (Laboratory Animal Science Association, 1998). Venous tail bleed samples were then taken at 2, 5, 10, 20, 40 and 60 min post-injection, keeping sampling volumes within LASA guidelines (Laboratory Animal Science Association, 1998). Mice were kept anaesthetised with isoflurane throughout the experiment. Blood was collected into Microvette 300μL lithium heparin capillary tubes (Sarstedt, Nümbrecht, Germany) and centrifuged at 1500x *g* for 10 min. Plasma supernatant was removed and stored at −80°C prior to analysis. Individual plasma samples were analysed following 1:9 plasma:deionised water.

### 2.6 LC-QTOF-MS analytical conditions

Analysis of plasma and urine was performed on an Agilent 1290 Infinity HPLC coupled to an Agilent 6550 QTOF-MS (Agilent). Data acquisition parameters are detailed in Appendix 1.

### 2.7 Design of LC-QTOF-MS profiling analyses

Samples from Experiments 1 (*Hgd*^−/−^ vs *Hgd*^+/−^) and 3 (^13^C_6_ HGA metabolic flux analysis) were analysed in separate batches, each comprising negative then positive polarity. The analytical sequence of each profiling batch was designed according to published guidance (Vorkas et al., 2015) and following the procedure described previously by Norman *et al*. (Norman et al., 2019).

### 2.8 Data processing and statistical analyses

Mining of metabolite features in raw data was performed using two parallel approaches (Figure 1, Experiment 1). A targeted approach was used to extract signals matching a 466-compound AMRT database previously generated in our laboratory from IROA Technology MS metabolite library of standards, accessible via https://doi.org/10.6084/m9.figshare.c.4378235.v2 (Norman et al., 2019), or a compound database comprising accurate masses of additional compounds with potential relevance to AKU. A complementary non-targeted approach was used to extract unknown chemical entities.

#### 2.8.1 Targeted feature extraction

Targeted feature extraction was performed in Profinder (build 08.00, Agilent) using the chemical formulae of compounds from the AMRT database described above. Extraction parameters were accurate mass match window ±10ppm and retention time (RT) window ±0.3 min. Allowed ion species were: H^+^, Na^+^, and NH_4_^+^ in positive polarity, and H^−^ and CHO_2_− in negative polarity. Charge state range was 1-2, and dimers were allowed. ‘Find by formula’ filters were: score >60 in at least 60% of samples in at least one sample group (samples were grouped by *Hgd*^−/−^ or *Hgd*^+/−^ and pre- or on nitisinone).

Eighty-five additional metabolites of potential interest in AKU or from wider tyrosine metabolism were appended to the database for targeted extraction. Forty-three were from the following pathway databases available from Pathways to PCDL (build 07.00, Agilent): ‘citrate degradation’, ‘noradrenaline and adrenaline degradation’ and ‘superpathway of phenylalanine, tyrosine and tryptophan biosynthesis’ (Appendix 2). Six metabolites were added as they were predicted to show potential alteration in *Hgd*^−/−^ due to association with tyrosine conjugation (acetyl-L-tyrosine, and γ-glutamyl-tyrosine) based on a previous publication (Norman et al., 2019) or association with ochronotic pigment derived from HGA (2,5-dihydroxybenzaldehyde, di-dehydro-homogentisic acid, hipposudoric acid and norhipposudoric acid). Thirty-six metabolites were from a list of potential biotransformation products directly derived from HGA and compiled using the Biotransformation Mass Defects application (Agilent; Appendix 3). This tool provides a list of potential metabolic biotransformation products covering both phase I (n=19) and II (n=17) metabolism for a given compound based on empirical formula. The data were mined for these additional metabolites with putative identification by accurate mass (±5ppm) only.

#### 2.8.2 Non-targeted feature extraction

Non-targeted extraction was performed by recursive feature extraction in Profinder (build 08.00). Extraction parameters are detailed in Appendix 4.

#### 2.8.3 Isotopologue feature extraction on data from ^13^C_6_ HGA metabolic flux analysis

Data from Experiment 3 (Figure 1) were mined using batch isotopologue extraction in Profinder (build 08.00). Here, compounds that showed significant differences between *Hgd*^−/−^ and *Hgd*^+/−^ mice (Experiment 1) were investigated for potential association with the ^13^C_6_ HGA tracer by examining the relative abundances of their M+0 to M+6 isotopologues. Extraction was performed with accurate mass and RT match windows of ±5ppm and ±0.3 min respectively against an AMRT database consisting only of these compound targets.

#### 2.8.4 Data QC and statistical analyses

First, initial QC was performed on all datasets in Profinder by manual curation of the dataset to remove visually low quality peaks and to correct integration issues across the dataset where appropriate.

Urine profiling datasets were then exported from Profinder as .csv files and imported into Mass Profiler Professional (build 14.5, Agilent) for additional QC and subsequent statistical analyses. First, creatinine peak area from this analysis creatinine was used as an external scalar for each mouse sample to correct for differences in urine concentration, as described previously (Norman et al., 2019). Additional QC was performed using data from pooled samples, which were interspersed throughout each analytical sequence. Entities were retained if a) observed in 100% of replicate injections for at least one sample group pool, and b) peak area coefficient of variation (CV) remained <25% across replicate injections for each sample group pool. Statistically significant entities were then identified in each dataset by t-tests; unpaired t-tests for *Hgd*^−/−^ vs *Hgd*^+/−^ comparisons, and paired t-tests for pre- vs on nitisinone comparisons. Multiple testing correction was performed using Benjamini-Hochberg false discovery rate (FDR) adjustment. FC’s were log_2_-transformed and based on peak area. FC cut-off was not applied for entities with FDR-adjusted p<0.05 however, in order to retain entities that showed consistent although relatively low magnitude abundance differences between comparison groups. Principal Components Analysis (PCA) was performed on each filtered dataset using four-component models.

The results from isotopologue extraction on plasma ^13^C_6_ HGA metabolic flux data were reviewed visually in Profinder for clear evidence of an isotope label likely to be derived from HGA. This data review was performed for compound matches individually, considering the predicted number of ^13^C atoms derived from the HGA tracer based on the compound’s chemical structure.

### 2.9 Study approval

All animal work was carried out in accordance with UK Home Office Guidelines under the Animals (Scientific Procedures) Act, 1986.

Metabolomic analyses on patient samples was approved by the Royal Liverpool and Broadgreen University Hospital Trust’s Audit Committee (audit no. ACO3836) and was part of the diagnostic service to patients being seen at the NAC.

## 3. Results

PCA showed clear separation in principal component 1 between the urine profiles of *Hgd*^−/−^ and *Hgd*^+/−^ mice from targeted (Figures 2A & 2B) and non-targeted feature extraction. Results from targeted and non-targeted extraction are presented separately in the following sections (number of entities obtained in each extraction method summarised in Appendix 5). Data from all animals in each experiment were included in the analysis.

**Figure 2.**
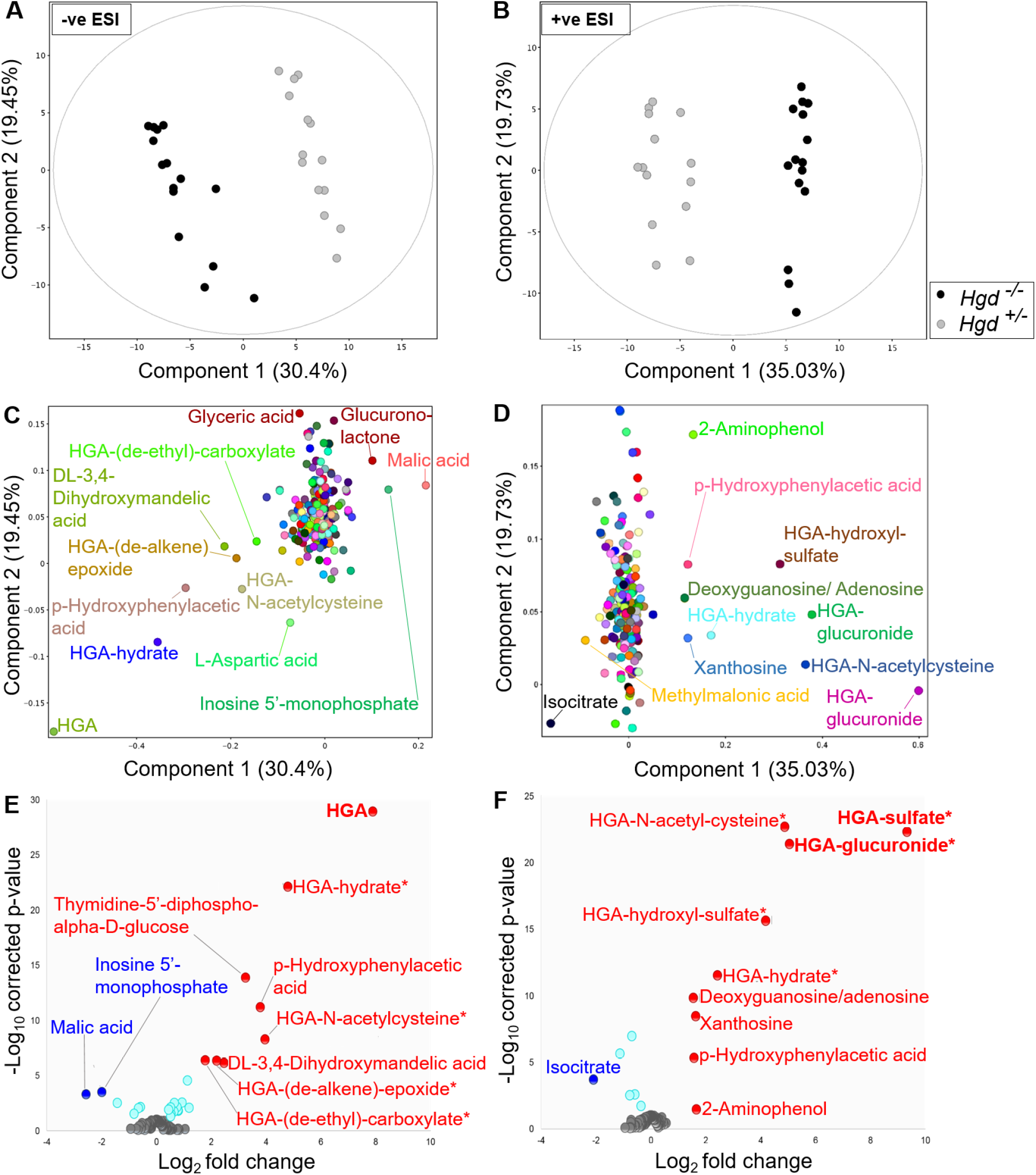
Clear differences between the urine metabolomes of *Hgd*^−/−^ and *Hgd*^+/−^ mice. A-D: PCA on data from targeted feature extraction, with PCA plots showing separation between *Hgd*^−/−^ and *Hgd*^+/−^ mice by component 1 in A, negative, and B, positive ionisation polarities. Lower plots show the corresponding PCA loadings of metabolites on components 1 and 2 in C, negative, and D, positive polarity. E-F: volcano plots illustrating selection of statistically significant urinary chemical entities between *Hgd*^−/−^ and *Hgd*^+/−^ mice based on p-value and fold change. E, negative polarity; F, positive polarity. Entities with p<0.05 (Benjamini-Hochberg FDR adjusted) and log_2_ fold change >1.5 are labelled, with red and blue indicating increased and decreased abundance, respectively, in *Hgd*^−/−^. Turquoise indicates adjusted p<0.05 but log_2_ fold change <1.5. Bold text indicates that the increase observed in *Hgd*^−/−^ was confirmed in mouse plasma following injection with ^13^C_6_ HGA tracer. * Compound not previously reported in the literature.

### 3.1 Targeted feature extraction

Targeted feature extraction was performed to search for metabolites based on AMRT (accurate mass ±10ppm, RT ±0.3 min) or accurate mass (±5ppm) alone. 27/250 and 15/243 metabolites showed abundance differences (FDR-adjusted p<0.05) between *Hgd*^−/−^ and *Hgd*^+/−^ in negative and positive polarity, respectively. Table 1 shows the altered metabolites ranked by FC. PCA loadings plots (Figures 2C & 2D) and volcano plots (Figures 2E & 2F) show that the greatest differences between *Hgd*^−/−^ and *Hgd*^+/−^ urine were in metabolites associated with HGA in negative and positive polarity. HGA and 8 predicted HGA biotransformation products were markedly elevated in *Hgd*^−/−^. Interestingly, HGA-sulfate (FC=9.3, p<0.0001) showed a greater FC increase than HGA (FC=7.9, p<0.0001). Six other HGA products were increased with FC >1.5 and p<0.0001: HGA-glucuronide, HGA-hydroxyl-sulfate, HGA-N-acetylcysteine, HGA-hydrate, HGA-(de-alkene)-epoxide and HGA-(de-ethyl)-carboxylate. Acetyl-HGA was elevated in *Hgd*^−/−^ (FC=2.3, p<0.0001), but did not pass QC filtering by CV <25% across replicate injections of pooled QC samples; a decrease in signal across the run indicated a stability issue (despite the auto-sampler being maintained at 4°C).

**Table 1.** Summary of identified metabolites or chemical entities showing altered abundance in *Hgd*^−/−^ mice. Direction of alteration and log_2_ fold change is indicated in *Hgd*−/− (relative to *Hgd*+/−) and following nitisinone treatment in *Hgd*−/− (paired data; on nitisinone relative to pre-nitisinone). P-values are false discovery rate adjusted. Where entities were significantly different in positive and negative polarity, the result with the lowest fold change is provided.

| Compound | Neutral mass | RT (min) | Direction of alteration in *Hgd<sup>-/-</sup>* | Log<sub>2</sub> fold change | p-value |
| --- | --- | --- | --- | --- | --- |
| HGA-sulfate | 247.9996 | 2.6 | ↑ | 9.3 | <0.0001 |
| HGA* | 168.0434 | 3.4 | ↑ | 7.9 | <0.0001 |
| HGA-glucuronide | 344.075 | 2.4 | ↑ | 5 | <0.0001 |
| HGA-hydroxyl-sulfate | 263.9931 | 2.8 | ↑ | 4.2 | <0.0001 |
| HGA-N-acetylcysteine | 329.057 | 4.5 | ↑ | 4 | <0.0001 |
| Thymidine-5'-diphospho-alpha-D-glucose* | 564.0758 | 2.4 | ↑ | 3.3 | <0.0001 |
| Unknown | 268.9662 | 2.6 | ↑ | 2.8 | <0.0001 |
| Unknown | 367.0289 | 3.4 | ↑ | 2.6 | <0.0001 |
| DL-3,4-Dihydroxymandelic acid* | 184.037 | 1.9 | ↑ | 2.5 | <0.0001 |
| HGA-hydrate | 186.053 | 3.4 | ↑ | 2.4 | <0.0001 |
| Unknown | 507.141 | 3.4 | ↑ | 2.4 | <0.0001 |
| Acetyl-HGA<sup>‡</sup> | 210.053 | 6.2 | ↑ | 2.3 | <0.0001 |
| Unknown | 287.9939 | 6.0 | ↑ | 2.3 | 0.025 |
| HGA-(de-alkene)-epoxide | 184.0369 | 1.8 | ↑ | 2.2 | <0.0001 |
| 2-Amino-6-(benzylthio)purine<sup>†</sup> | 257.0758 | 3.5 | ↑ | 2.2 | <0.0001 |
| Unknown | 257.0391 | 3.4 | ↑ | 2.0 | <0.0001 |
| HGA-(de-ethyl)-carboxylate | 184.0017 | 3.4 | ↑ | 1.8 | <0.0001 |
| 2-Aminophenol* | 109.0534 | 1.9 | ↑ | 1.7 | 0.035 |
| Unknown | 294.0949 | 3.6 | ↑ | 1.7 | <0.001 |
| p-Hydroxyphenylacetic acid* | 152.0478 | 5.5 | ↑ | 1.6 | <0.0001 |
| Unknown | 361.0827 | 1.7 | ↑ | 1.5 | <0.001 |
| 3-Dehydroshikimic acid* | 172.0373 | 1.8 | ↑ | 1.2 | <0.01 |
| Hydroxymethyl-HGA | 198.0526 | 4.8 | ↑ | 1.1 | <0.0001 |
| Aceturic acid* | 117.0426 | 1.7 | ↑ | 0.94 | <0.01 |
| Nicotinic acid* | 123.032 | 1.8 | ↑ | 0.8 | 0.015 |
| 4-Hydroxybenzaldehyde* | 122.0368 | 5.6 | ↑ | 0.8 | 0.014 |
| Vanillylmandelic acid* | 198.0528 | 3.3 | ↑ | 0.75 | <0.01 |
| N-Acetyl-DL-serine* | 147.0532 | 1.6 | ↑ | 0.66 | <0.01 |
| N-Acetylaspartate* | 175.0481 | 1.8 | ↑ | 0.64 | 0.018 |
| Xanthosine* | 284.0762 | 3.4 | ↑ | 0.53 | 0.028 |
| N-Alpha-acetyl-l-asparagine* | 174.0641 | 1.6 | ↑ | 0.52 | <0.01 |
| N-Acetyl-L-phenylalanine* | 207.0895 | 6.8 | ↑ | 0.52 | 0.049 |
| 3,5-Cyclic-AMP* | 329.0525 | 3.4 | ↑ | 0.44 | 0.012 |
| Malic acid* | 134.0211 | 1.5 | ↓ | 2.6 | <0.001 |
| Isocitrate* | 192.0278 | 1.7 | ↓ | 2.1 | <0.001 |
| Vanillyl pyruvic acid† | 256.0582 | 5.0 | ↓ | 2.1 | <0.001 |
| Inosine 5'-monophosphate* | 348.05 | 1.8 | ↓ | 2 | <0.001 |
| Unknown | 242.0427 | 4.0 | ↓ | 1.7 | <0.0001 |
| Unknown | 362.086 | 9.6 | ↓ | 1.6 | <0.01 |
| Glucuronolactone* | 176.0321 | 1.6 | ↓ | 1.4 | <0.01 |
| Methylmalonic acid* | 118.0266 | 2.0 | ↓ | 1.1 | <0.0001 |
| Monomethyl-glutaric acid* | 146.0579 | 5.1 | ↓ | 0.82 | 0.029 |
| N-Acetyl-L-glutamic acid* | 189.0637 | 2.1 | ↓ | 0.76 | <0.01 |
| Citramalic acid* | 148.039 | 2.6 | ↓ | 0.7 | <0.0001 |
| L-anserine* | 240.1222 | 1.3 | ↓ | 0.59 | <0.01 |
| 2,3-Cyclic AMP* | 329.0525 | 2.7 | ↓ | 0.58 | <0.01 |
| O-Succinyl-L-homoserine* | 219.0743 | 2.0 | ↓ | 0.35 | 0.019 |

Excluding HGA, 26 significantly altered entities, were AMRT-matched with compounds from the 466-compound library developed in-house. The most significantly altered (p<0.05 and log_2_ FC>1.5) of these were: thymidine-5’-diphospho-alpha-D-glucose, DL-3,4-dihydroxymandelic acid, 2-aminophenol, p-hydroxyphenylacetic acid (increased in *Hgd*^−/−^), malic acid, isocitrate and inosine 5’-monophosphate (decreased in *Hgd*^−/−^).

### 3.2 Non-targeted feature extraction

Non-targeted feature extraction yielded 359 and 213 chemical entities post-QC in negative and positive polarity respectively; mass range = 54-3108 Da and RT range = 1.1-11.5 min. Comparison of *Hgd*^−/−^ and *Hgd*^+/−^ revealed entities with clear abundance differences (p<0.05, log_2_ FC>1.5); 9 in negative polarity (6 increased in *Hgd*^−/−^, 3 decreased in *Hgd*^−/−^) and 2 in positive polarity (both increased in *Hgd*^−/−^). The mass range of these significant entities was 242-507 Da, and RT range was 1.7-9.6 min (Table 1). Candidates compound matches were obtained for two of these entities based on accurate mass match (<10ppm) with the MassHunter METLIN metabolite PCD/PCDL accurate mass database (build 07.00; accessed through Agilent PCDL Manager build 8.00); 2-amino-6-(benzylthio)purine (increased in *Hgd*^−/−^) and vanilyl pyruvic acid (decreased in *Hgd*^−/−^).

### 3.3 The effect of nitisinone treatment on chemical entities altered in *Hgd*^−/−^ mice

To investigate the effect of nitisinone treatment on the entities shown to be altered here in *Hgd*^−/−^, they were searched in the data from previous mouse and human urine profiling experiments (Figure 1, Experiment 2). These data were acquired from *Hgd*^−/−^ mice at baseline then after 1 week on nitisinone, and from patients with AKU at baseline then after 24 months on nitisinone (Norman et al., 2019).

Eight entities from targeted feature extraction that were altered in *Hgd*^−/−^ vs *Hgd*^+/−^ mice were significantly altered in the opposite direction in urine from both *Hgd*^−/−^ mice and patients with AKU on nitisinone (based on peak area pre- vs on nitisinone, Benjamini-Hochberg FDR-adjusted p<0.05; Table 2); 7 decreased, and 1 increased. The 7 decreased metabolites were HGA, the HGA biotransformation products HGA-sulfate, HGA-glucuronide, HGA-hydrate and hydroxymethyl-HGA, and also xanthosine and 3,5-cyclic-AMP. The increased metabolite was monomethyl-glutaric acid. Interestingly, p-hydroxyphenylacetic acid and 4-hydroxybenzaldehyde were increased both in *Hgd*^−/−^ vs *Hgd*^+/−^ and also on nitisinone in *Hgd*^−/−^ mice and patients.

**Table 2.**
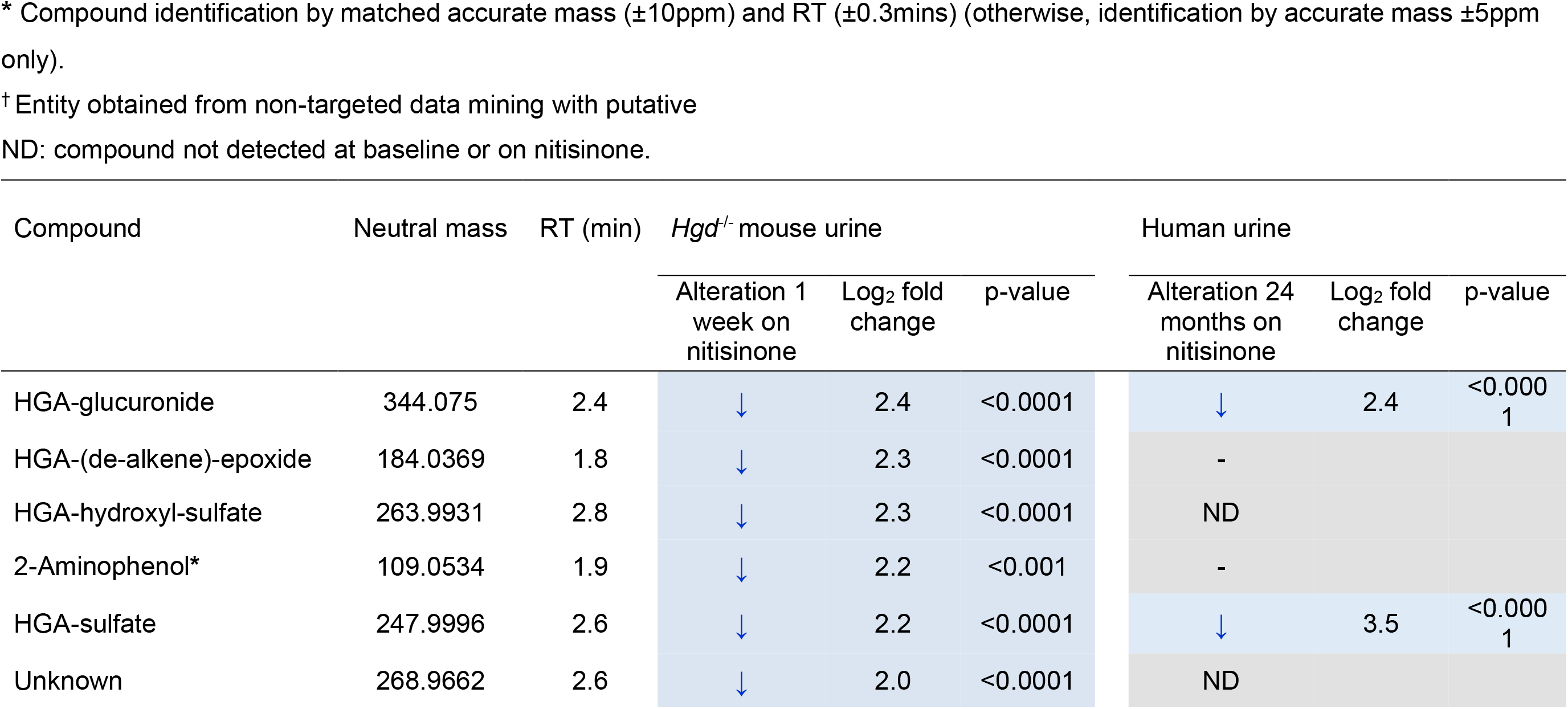
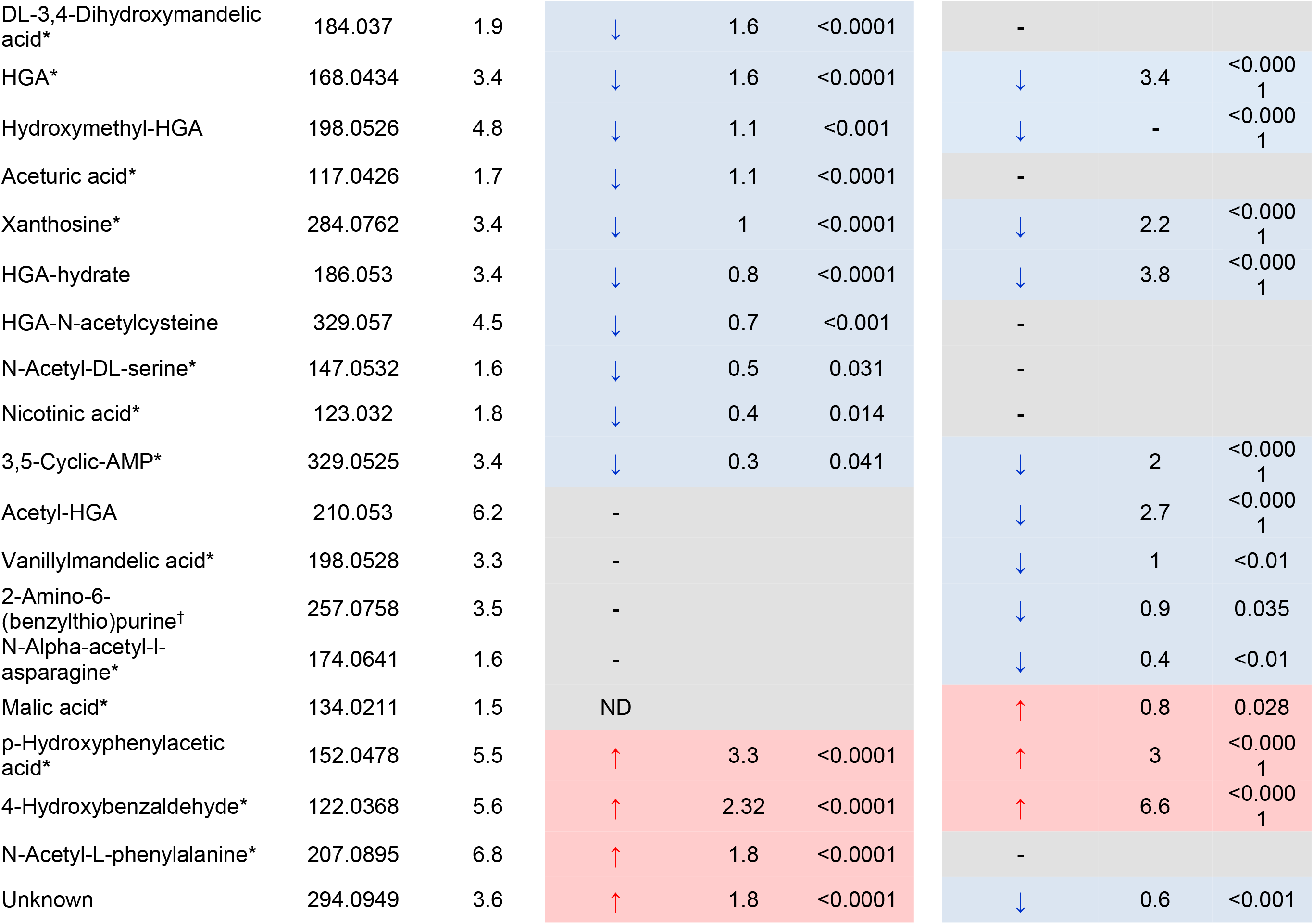
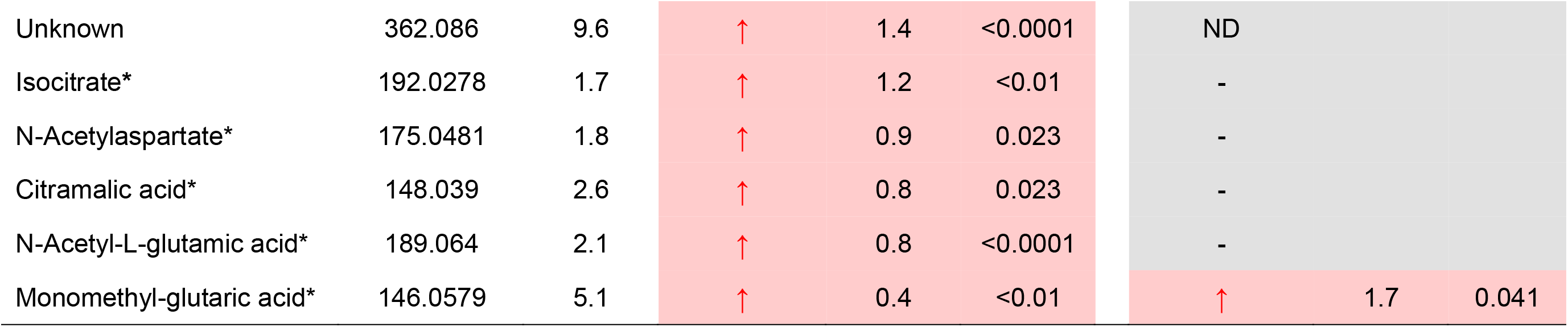
The effect of nitisinone treatment in *Hgd*^−/−^ mice and patients with AKU on the abundance of urinary chemical entities altered in *Hgd*^−/−^ vs *Hgd*^+/−^ mice. The entities that showed differences between *Hgd*^−/−^ vs *Hgd*^+/−^ mice (Table 1) were examined in two additional datasets. Paired t-tests were employed to compare the abundances at baseline versus 1 week on nitisinone (4mg/L, in drinking water) for *Hgd*^−/−^ mice and 24 months on 2mg daily nitisinone for patients with AKU. Only entities with false discovery rate adjusted p<0.05 pre- vs on nitisinone in mouse or human are displayed. Direction of alteration and log_2_ fold change are indicated. Where entities were significantly different in positive and negative polarity, the result with the lowest fold change is provided. Note: no fold change indicated for hydroxymethyl-HGA in humans, as this compound was not detected for any patient on nitisinone.

Nineteen entities that were altered in *Hgd*^−/−^ vs *Hgd*^+/−^ mice were significantly altered in the opposite direction on nitisinone in urine from either *Hgd*^−/−^ mice (n=14; 9 decreased, 5 increased on nitisinone) or patients (n=5; 4 decreased, 1 increased on nitisinone) only (Table 2). These compounds comprised the remaining HGA biotransformation products, except HGA-(de-ethyl)-carboxylate, which were all decreased on nitisinone. On nitisinone, acetyl-HGA was decreased in patients only, and HGA-hydroxyl-sulfate, HGA-N-acetylcysteine and HGA-(de-alkene)-epoxide were decreased in mice only.

### 3.4 Confirmation of HGA biotransformation products by ^13^C_6_ HGA metabolic flux analysis (Experiment 3)

Data from isotopologue extraction were compared between plasma collected from the same mice across the time intervals available (2-60 min). The M+6 isotopologue was of particular interest as mice were injected with ^13^C_6_-labelled HGA (although M+ 0-5 isotopologues were also considered for compounds where the labelled ^13^C carbons of the benzene ring could potentially be removed through metabolism of HGA).

A clear M+6 peak for HGA was observed over the sampling time course in plasma from *Hgd*^−/−^ and *Hgd*^+/−^ mice, although the signal decreased from 10-20 min post-injection (Figure 3). As indicated in Figure 2 (compounds in bold text), 2 HGA biotransformation products that were increased in *Hgd*^−/−^ urine were observed with clear M+6 peaks over the time course; HGA-glucuronide and HGA-sulfate. These data confirm that the compounds are derived from HGA. HGA-sulfate showed a similar time course profile to HGA. For this compound the native M+0 isotopologues were also absent from *Hgd*^+/−^ plasma at all time points, in contrast to the M+6 isotopologue whose profile appeared to closely follow that of the HGA M+6 peak over the time course in both *Hgd*^−/−^ and *Hgd*^+/−^ (Figure 3). For HGA-glucuronide, the native M+0 isotopologue was also absent from *Hgd*^+/−^ plasma, but the M+6 isotopologue was only observed in *Hgd*^−/−^ (Figure 3), indicating that glucuronidation of the HGA tracer was evident only for *Hgd*^−/−^ mice.

**Figure 3.**
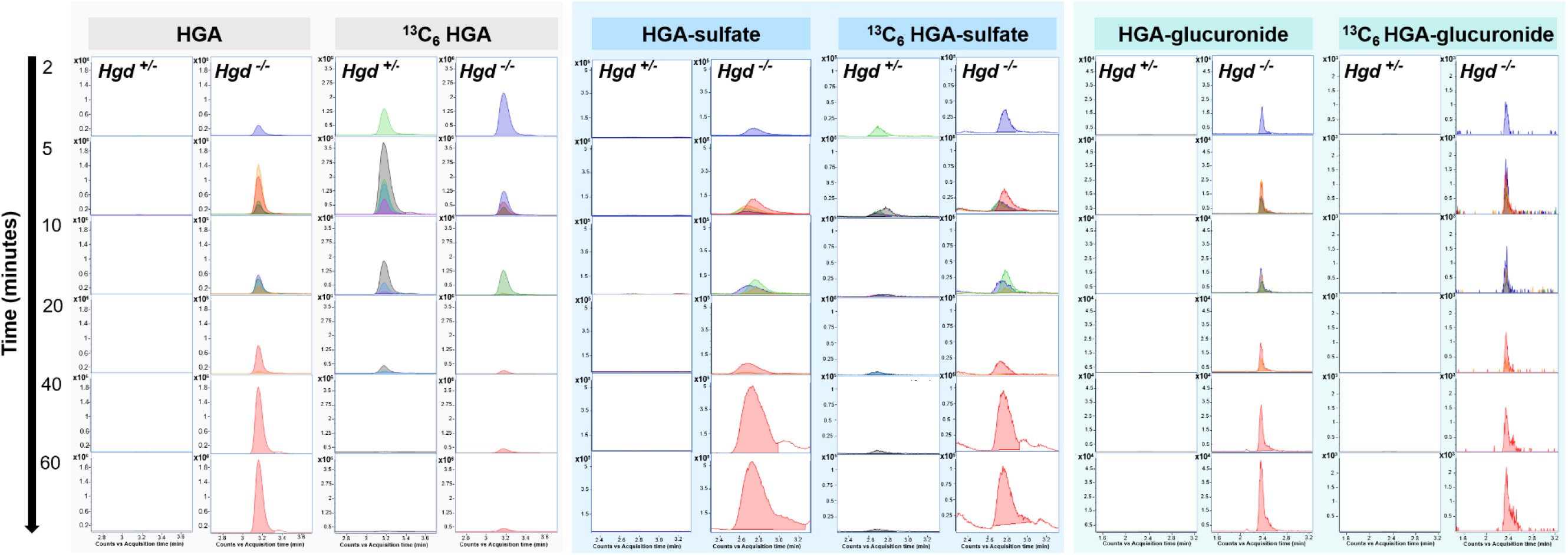
Isotopologue extraction results on plasma from the *in vivo* metabolic flux experiment using injected ^13^C_6_-labelled homogentisic acid (HGA). Data shown are from *Hgd*^−/−^ and *Hgd*^+/−^ samples taken at intervals of 2, 5, 10, 20, 40 and 60 (when possible) min after injection. Extracted ion chromatograms (EIC’s) represent the M+0 (native compound) and M+6 (^13^C_6_-labelled form) isotopologue signals for HGA, HGA-sulfate and HGA-glucuronide. EIC’s show clear M+6 peaks for these compounds following injection (but only from *Hgd*^−/−^ mice for HGA-glucuronide), confirming that they are derived from the labelled HGA.

## 4. Discussion

The data reported here from the first metabolome-wide comparison of non-treated AKU vs non-AKU (*Hgd*^−/−^ vs *Hgd*^+/−^ mice; Experiment 1) show that the biochemical consequences of HGD-deficiency are wider-reaching than previously recognised and extend beyond tyrosine metabolism. The data further reveal for the first time that phase I and II metabolic processes are recruited for detoxification of the markedly increased HGA. The observation of a reversed direction of alteration for a number of these urinary metabolites following nitisinone treatment in mice and patients (Experiment 2) indicates that these changes observed in the mouse model of AKU apply to the human disease.

### 4.1 HGA biotransformation products from phases I and II metabolism

The clearest differences between the urine metabolomes of *Hgd*^−/−^ versus *Hgd*^+/−^ mice were associated with HGA; HGA and 8 previously unreported HGA-derived biotransformation products were increased in *Hgd*^−/−^. Figure 4 shows the predicted structures for 7 of these novel compounds and the enzyme families responsible for their production. Reversed alteration (decreases) in HGA-sulfate, HGA-glucuronide, HGA-hydrate, acetyl-HGA and hydroxymethyl-HGA for patients on nitisinone (Experiment 2) adds validation and indicates that these HGA biotransformation processes are recruited in human AKU.

**Figure 4.**
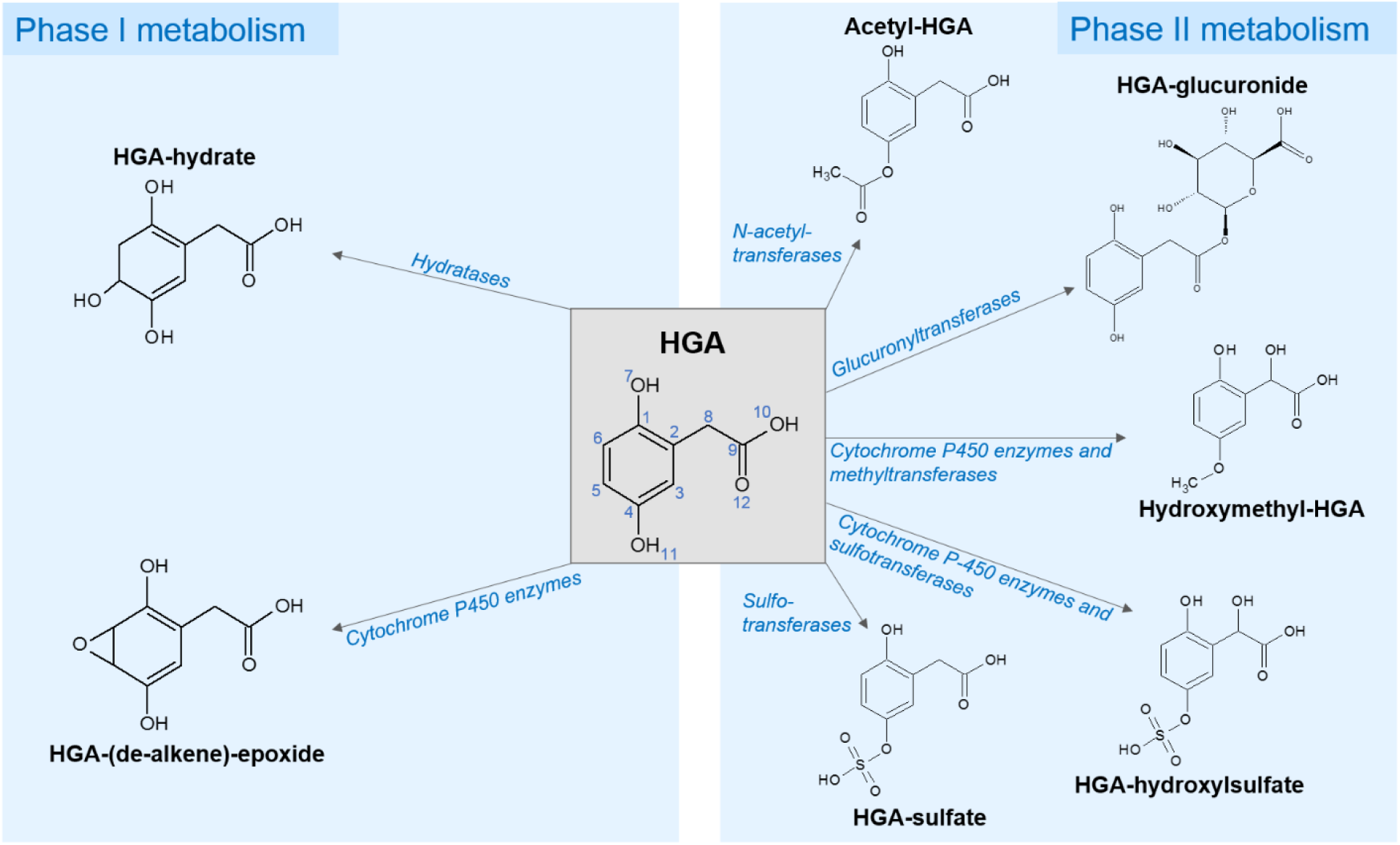
Predicted structures of newly-identified HGA clearance products resulting from phase I and II metabolism. The proposed sites for metabolism/conjugation were those with the greatest probability from *in silico* prediction tools Stardrop (Optibrium, Cambridge, UK) and Reacting Atom (Way2Drug, Moscow, Russia), for phase I and II biotransformations respectively. Note: HGA positions labelled 7 and 11 have equal probability for sulfation (deltaP = 0.8).

Observation of ^13^C_6_-labelled forms of HGA-glucuronide and HGA-sulfate in plasma following ^13^C_6_-HGA injection in mice (Experiment 3) confirmed that these products are derived from HGA. ^13^C_6_-HGA-sulfate was observed in *Hgd*^−/−^ and *Hgd*^+/−^ mice, following the profile of the ^13^C_6_-HGA across the sampling time course. Interestingly, ^13^C_6_-HGA-glucuronide was only observed in *Hgd*^−/−^, suggesting prior upregulation of glucuronyltransferase activity for HGA clearance in AKU. HGA-glucuronide (FC=5), HGA-sulfate (FC=9.3) and HGA (FC=7.9) showed the greatest increases in *Hgd*^−/−^ versus *Hgd*^+/−^.

To our knowledge, these are the first data on HGA metabolism outside of its conversion to maleylacetoacetic acid in the traditional tyrosine degradation pathway. Phase I and II pathways are traditionally considered as routes for drug metabolism with their enzymes primarily located in the liver; as with the enzymes of the tyrosine pathway (Williams et al., 2012). Phase I metabolism comprises reactions to introduce reactive or polar groups to substrates and includes hydroxylation, oxidation, reduction and hydrolysis. Phase II metabolism involves conjugation reactions to form glucuronide, glutathione, mercapturic acid, amino acid, methyl and acetyl conjugates (Williams, 1971; Wolff and Jordan, 1987). Phase I and II reactions can also occur sequentially; here the compounds hydroxymethyl-HGA and HGA-hydroxylsulfate were formed by hydroxylation with subsequent methylation (hydroxymethyl-HGA) or sulfation (HGA-hydroxylsulfate). Not all phase I and II biotransformations are adaptive clearance mechanisms. HGA-(de-alkene)-epoxide is a product of HGA bioactivation; epoxide metabolites are generally reactive, electrophilic compounds that require further enzymatically-mediated detoxification (Obach and Kalgutkar, 2010; Hughes et al., 2015b). It is also worth noting the potential HGA biotransformation products that were not detected in *Hgd*^−/−^ in this study (see Appendix 3 for the full list of potential HGA-derived products predicted *in silico*). A particularly notable example is the absence of a glycine conjugate of HGA, as glycine conjugation is a known detoxification mechanism of a number of other aromatic acids (Knights et al., 2007; Knights and Miners, 2012).

The magnitude of these alterations is of potential clinical importance in AKU as they reveal for the first time the specific metabolic processes that are major detoxification mechanisms to render HGA chemically inert. HGA is the primary toxic agent in AKU; its oxidation produces ochronotic pigment, which becomes bound within the extracellular matrix of collagenous tissues. The benzoquinone intermediates in ochronotic pigment formation are highly reactive species capable of inducing oxidative changes to proteins and lipids (Hegedus and Nayak, 1994; Braconi et al., 2015). These oxidative alterations are likely to self-perpetuate osteoarthropathy associated with ochronosis and could account for cases of acute fatal metabolic consequences reported in the literature (Davison et al., 2016). Identification of the specific enzymes that catalyse these HGA biotransformations could inform future AKU therapeutic interventions aimed at enhancing HGA degradation. Glucuronidation, for example, is quantitatively one of the most important phase II biotransformation reactions, carried out by 15 UPD-glucuronosyltransferase enzymes in humans (Burchell et al., 1991; Tukey and Strassburg, 2000). A number of agents, including naturally occurring dietary compounds, are known to be potent inducers of UPD-glucuronosyltransferases and other phase II enzymes (Talalay et al., 1988; Prestera et al., 1993; Fahey and Talalay, 1999); our data show that such agents have potential use in diseases caused by toxic metabolite accumulation, including inborn errors of metabolism.

### 4.2 Associated alteration to tyrosine, purine and TCA cycle metabolism in AKU

This study showed for the first time that in untreated AKU there is alteration to tyrosine, purine and TCA cycle metabolites (Figure 5). These data add to a growing body of evidence from metabolomic studies that perturbations in tyrosine, purine and energy metabolism are linked in numerous diseases including Alzheimer’s (Kaddurah-Daouk et al., 2011, 2013), depression (Weng et al., 2015) and diabetic nephropathy (Men et al., 2017).

**Figure 5.**
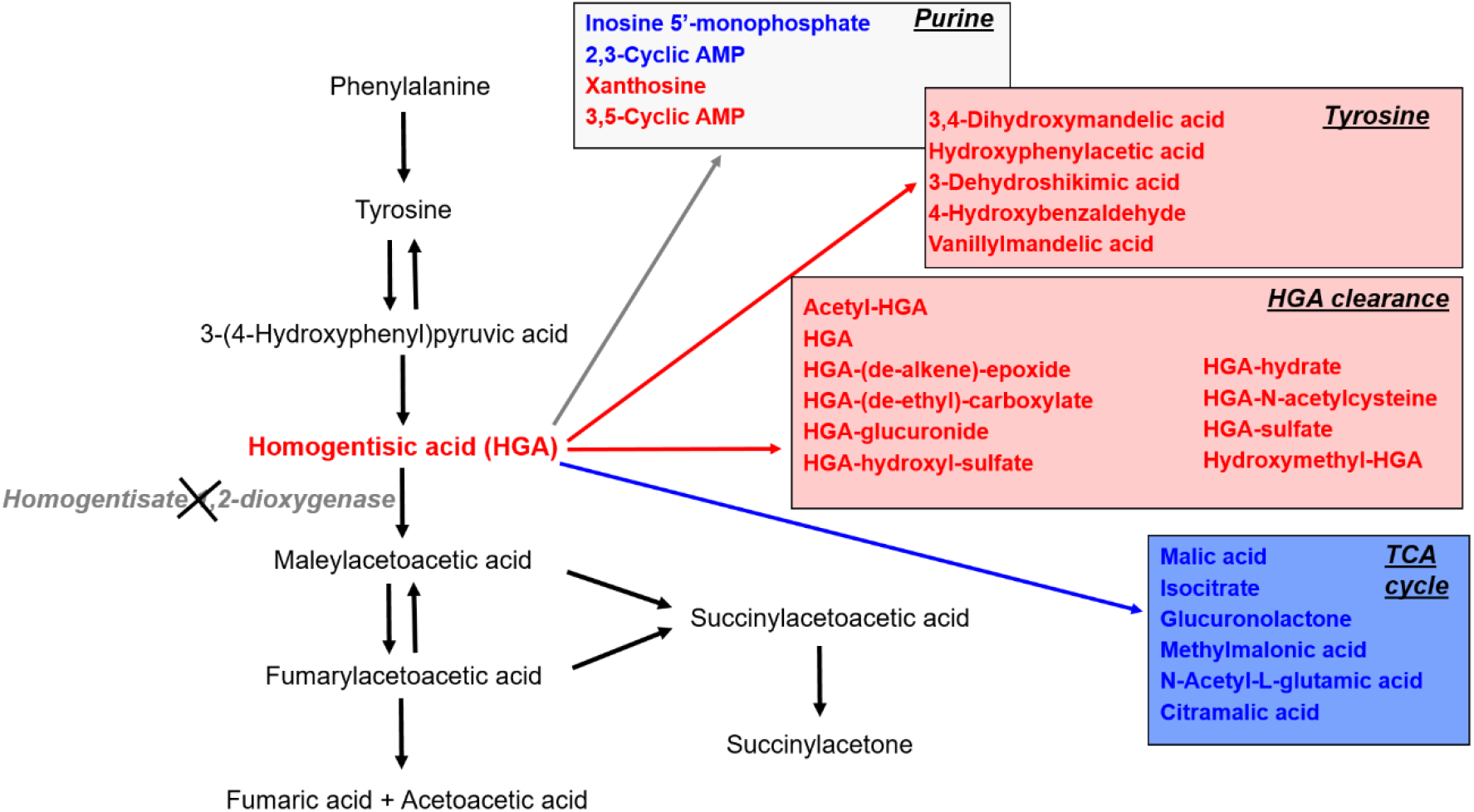
Summary of metabolites altered in *Hgd*^−/−^ mouse urine grouped by their associated pathways. Left: the tyrosine degradation pathway showing lack of the enzyme HGD in AKU and the consequential increase in HGA. Right (boxes): observed metabolite alterations grouped by pathway; red and blue indicate increased and decreased abundance respectively. Tyrosine metabolites including HGA, and HGA clearance products from phase I and II metabolism were elevated. Metabolites associated with the TCA cycle were decreased. A combination of increased and decreased abundance was observed for purine pathway metabolites.

Previous metabolomic studies in AKU have focused solely on the impact of nitisinone on the metabolome (Davison et al., 2018a, 2018b, 2019a, 2019b; Norman et al., 2019). Nitisinone reversibly inhibits hydroxyphenylpyruvic acid dioxygenase (HPPD; E.C. 1.13.11.27), the enzyme that produces HGA, and it is currently the most effective treatment for AKU. Nitisinone reduces plasma and urine HGA concentrations (Phornphutkul et al., 2002; Introne et al., 2011; Olsson et al., 2015; Ranganath et al., 2016; Milan et al., 2017), completely arrests ochronosis in AKU mice (Preston et al., 2014; Keenan et al., 2015) and more recently was shown to slow progression of morbidity in patients attending the UK NAC (2mg nitisinone daily) (Ranganath et al., 2018). We showed previously in serum and urine that nitisinone induced an extended network of metabolic alteration within tyrosine and neighbouring pathways, including tryptophan, purine and TCA cycle (Davison et al., 2019a; Norman et al., 2019). This alteration is a concern in AKU, and particularly in hereditary tyrosinaemia type-1, another inherited disorder of tyrosine metabolism, in which nitisinone treatment is essential from early infancy (McKiernan et al., 2015).

#### 4.2.1 Alteration to tyrosine metabolism

The increases in tyrosine metabolites, excluding HGA, were unexpected in untreated AKU. Increases in metabolites upstream of HPPD previously reported in nitisinone-treated AKU were thought to be a direct consequence of the inhibition of HPPD by nitisinone and the consequential hypertyrosinaemia (Davison et al., 2019a; Norman et al., 2019). For the first time, the present data show that targeted *Hgd* disruption in *Hgd*^−/−^ mice induces metabolic changes upstream of HGA, despite no increase in tyrosine, HPPA or HPLA. The altered tyrosine metabolites were increased 3-dehydroshikimic acid, 3,4-dihydroxymandelic acid (DHMA),4-hydroxybenzaldehyde, hydroxyphenylacetic acid and vanillymandelic acid (VMA).

DHMA and VMA are catecholamine metabolites derived from noradrenaline. Noradrenaline is derived from tyrosine, via L-DOPA and dopamine, and metabolised to DHMA (via 3,4-dihydroxymandelaldehyde) then to VMA, or alternatively to adrenaline or normetadrenaline. It is surprising that these increases were observed in untreated AKU despite no increase in tyrosine. However, increased DHMA and VMA indicate increased metabolism of noradrenaline via 3,4-dihydroxymandelaldehyde as opposed to adrenaline or normetadrenaline in untreated AKU, which is supported by a previous observation of decreased urinary normetadrenaline in patients with AKU taking nitisinone (Davison et al., 2018a).

The cause of these unexpected changes to tyrosine and peripheral neurotransmitter metabolism is not clear. At the supraphysiological concentrations observed in AKU, HGA could potentially act on other enzymes of neurotransmitter metabolites and alter their activity. Alternatively, the changes could relate to an unknown feature of the disease. It is important to determine whether the changes observed in peripheral neurotransmitter metabolism indicate dysregulated central nervous system (CNS) homeostasis. Recent mass spectrometric mouse brain imaging analyses support that in nitisinone-treated AKU, CNS metabolic alteration is limited to tyrosine and tyramine (32). Further analyses of CNS tissues and biofluids (CSF) are required to provide a clearer picture of neurotransmitter metabolism in untreated AKU.

The finding that p-hydroxyphenylacetic acid and 4-hydroxybenzaldehyde were increased in *Hgd*^−/−^ mice and further increased on nitisinone is interesting. P-Hydroxyphenylacetic acid is generated from gut microbiota and its increase on nitisinone was thought to reflect enhanced microbial metabolism due to reduced oxidative stress upon commencement of treatment (Davison et al., 2019a). 4-Hydroxybenzaldehyde is a naturally occurring metabolite with therapeutic properties in numerous diseases (Kang et al., 2017). Benzaldehydes may relate to the proposed benzoquinone intermediates involved in the formation of ochronotic pigment from HGA, but the reason for the further increase in 4-hydroxybenzaldehyde following nitisinone is inconsistent with this and requires explanation.

#### 4.2.2 Alteration to purine metabolism

In nitisinone-treated AKU the purine metabolite changes previously reported were mainly decreased concentrations; 3-ureidoproprionate, adenine and allantoin were decreased in human urine, and 3,5-cyclic AMP and xanthosine were decreased in human and mouse urine (Norman et al., 2019). The present data indicate a more complex pattern of purine alteration in untreated AKU; increased 3,5-cyclic AMP and xanthosine, and decreased 2,3-cyclic AMP and inosine 5’-monophosphate.

Purine catabolism is important in the homeostatic response to various states of mitochondrial oxidative stress, with shifts occurring to favour breakdown to xanthine and uric acid (Kristal et al., 1999; Yao et al., 2010). The increased xanthosine and decreased 2,3-cyclic AMP and inosine 5’-monophosphate reported here for *Hgd*^−/−^ mice is consistent with a shift in purine catabolism in response to oxidative conditions induced by HGA and ochronosis, as xanthosine is a direct precursor for xanthine and uric acid. Xanthosine is increased in urine from rats with other oxidative diseases, including chronic kidney disease (Zhang et al., 2015) and diabetes nephropathy, and markedly decreased following administration of nephroprotective therapy (Men et al., 2017). The alteration in purine metabolism perhaps most associated with pathology is elevated uric acid, the classic example being gout, in which elevated blood uric acid is the direct cause of a painful arthropathy.

#### 4.2.3 Alteration to TCA cycle metabolism

Decreases in TCA-related metabolites in *Hgd*^−/−^ is the first indication of perturbed energy metabolism in untreated AKU. Decreases in citramalic acid, isocitrate and N-acetyl-L-glutamic acid were reversed on nitisinone in *Hgd*^−/−^, suggesting for the first time that nitisinone is at least partially restorative. Malic acid and isocitrate are major intermediates of the TCA cycle; the primary energy-producing pathway of the cell. Blood and urine levels of TCA metabolites generally reflect overall TCA cycle activity (Hallan et al., 2017). Inhibited TCA cycle activity could be due to overall mitochondrial biogenesis, decreased expression of genes encoding TCA cycle enzymes or reduced substrate availability; the latter seems more likely in AKU, in which HGD deficiency prevents further metabolism of HGA to the TCA cycle intermediate fumaric acid. Decreased bioavailability of fumaric acid may then explain the decreases observed for other TCA cycle metabolites. Previous analyses have investigated serum TCA cycle metabolites in AKU, but these were limited to studying the effect of nitisinone. In contrast to the present urinary data, the TCA metabolites succinic acid and α-ketoglutaric acid were decreased in serum from patients with AKU following nitisinone (Davison et al., 2019a).

Decreased urinary citrate (precursor for and structural isomer of isocitrate) is a well-known risk factor for kidney stone formation. It is possible that the decreased isocitrate reported here for *Hgd*^−/−^ mice is associated with the increased risk of kidney stones in AKU (Wolff et al., 2015). In this analysis isocitrate was the closest match obtained for this particular entity (RT=1.7 min) based on its AMRT in our database generated from standards (RT=1.6 min), but citric acid is also a potential match, with the same accurate mass but slightly different database RT (1.9 min). It is important to note that the database RTs were obtained from standards analysed in blank matrix, therefore a degree of RT shift can be expected in urine matrix due to effects such as pH. Hypocitratururia is generally defined as citric acid excretion <320mg (1.67mmol) per day in adults (Zuckerman and Assimos, 2009). The present profiling data are semi-quantitative; targeted quantitative assays are required to determine exact urinary citric acid concentrations in AKU patients and the additional therapeutic value of nitisinone in this regard.

## 5. Conclusions

In conclusion, we have compared, for the first time, the metabolome-wide profiles of untreated AKU (*Hgd*^−/−^ mice) versus non-AKU (*Hgd*^+/−^ mice). The data indicate that targeted homozygous disruption to the *Hgd* gene induces previously unknown metabolite changes in a complex network of pathways associated with altered tyrosine metabolism. Single-gene disorders, such as AKU, therefore present unique windows into metabolism that can uncover associations between metabolite pathways in physiology more generally. It was also revealed for the first time that a range of phase I and II metabolic processes are recruited for clearance of elevated HGA, which is central to the pathological features of AKU. These data have wider, clinically-significant implications beyond AKU by showing that phase I and II metabolic processes are recruited for detoxification of endogenous metabolites that can accumulate as a consequence of inherited metabolic disease. Our findings therefore highlight that it is a fundamental misconception to view these ancient, evolved processes solely as mechanisms for drug clearance. The wider take-home message is that rare diseases offer unique insights into metabolism, and even nature, more generally. As stated by the great physician, William Harvey (1578-1657); “nature is nowhere accustomed more openly to display her secret mysteries than in cases where she shows traces of her workings apart from the beaten path” (Wills, 1847).

## Abbreviations

AKU: alkaptonuria
HGD: homogentisate 1,2-dioxygenase
HGA: homogentisic acid
LC-QTOF-MS: liquid chromatography quadrupole time-of-flight mass spectrometry
AMRT: accurate mass / retention time
PCA: principal components analysis
RT: retention time
FDR: false-discovery rate
FC: fold change
QC: quality control
CV: coefficient of variation
CSF: cerebrospinal fluid
DHMA: DL-3,4-dihydroxymandelic acid
VMA: vanillylmandelic acid
CNS: central nervous system

## Acknowledgements

BPN is funded by the University of Liverpool, Royal Liverpool University Hospitals Trust and Agilent Technologies UK Ltd. ASD is funded through a National Institute for Health Research (NIHR) Doctoral Research Fellowship (grant code: HCS DRF-2014-05-009). JHH is funded by the Alkaptonuria Society. This article presents independent research partially funded by the NIHR. The views expressed are those of the author(s) and not necessarily those of the NHS, the NIHR or the Department of Health and Social Care. The authors thank Gordon A Ross and Hania Khoury of Agilent Technologies UK Ltd. for their guidance in isotopologue data extraction and review.

## Author contributions

**BPN, ASD:** Conceptualization, Methodology, Formal analysis, Investigation, Data curation, Writing – Original draft preparation, Writing – Review and Editing, Visualization. **JHH, HS, PJW, NGB, ATH, AMM:** Investigation, Writing – Review and Editing. **JCJ:** Resources, Writing – Review & Editing. **GBG:** Resources, Writing – Review and Editing, Supervision. **NBR, LRR, JAG:** Supervision, Writing – Review and Editing.

## Notes

### Competing Interest Statement

The authors have declared no competing interest.

