## Appendix 1 for "Studies in alkaptonuria reveal new roles beyond drug clearance for phase I and II biotransformations in tyrosine metabolism"

**Appendix 1. Quadrupole time-of-flight mass spectrometry conditions.**

Analysis of plasma and urine was performed on an Agilent 1290 Infinity LC coupled to an Agilent 6550 QTOF-MS equipped with a dual AJS electrospray ionization source (Agilent, UK). Reversed-phase HPLC was performed on an Atlantis dC_18_ column (3x100mm, 3µm, Waters, UK) maintained at 60°C. Mobile phase composition was (A) water and (B) methanol, both with 5mmol/L ammonium formate and 0.1% formic acid. The elution gradient began at 5% B 0-1 min and increased linearly to 100% B by 12 min, held at 100% B until 14 min, then at 5% B for a further 5 min. Sample injection volume was 2µL for all samples and the needle was washed with a solution of water:methanol:isopropanol (45:45:10 v/v) between injections. The autosampler compartment was maintained at 4°C.

The mass spectrometer was tuned and calibrated according to protocols recommended by the manufacturer. Acquisition was performed in 2 GHz mode, positive and negative ionisation polarity and mass range 50-1700. The capillary voltage was 4000 V and fragmentor voltage 380 V. The desolvation gas temperature was 200 °C with flow rate at 15 L/min. The sheath gas temperature was 300 °C with flow rate at 12 L/min. The nebulizer pressure was 40 psig and nozzle voltage 1000 V (± for positive and negative ionisation modes). The acquisition rate was 3 spectra/second.

A reference mass correction solution was prepared in 95:5 methanol:water containing 60 mg/dL (5 mmol/L) purine (C_5_H_4_N_4_, CAS No.: 120-73-0), 1310.4 mg/dL (100 mmol/L) trifluoroacetic acid ammonium salt (TFA; CF_3_CO_2_NH_4_, CAS No.: 3336-58-1) and 230.3 mg/dL (2.5 mmol/L) hexakis(1H, 1H, 3H-tetrafluoropropoxy)phosphazine (HP-0921; C_18_H_18_F_24_N_3_O_6_P_3_, CAS No.: 58943-98-9) (Agilent). The solution was continually infused at a flow rate of 0.5 mL/min by a separate isocratic pump for constant mass correction [positive ionisation: purine (m/z 121.0509), HP-0921 (m/z 922.0098); negative ionisation: TFA (m/z 112.9856), purine (m/z 119.0363), HP-0921 (HP-0921 + formate adduct: m/z 966.0007)].
