## Appendix 2 for "Studies in alkaptonuria reveal new roles beyond drug clearance for phase I and II biotransformations in tyrosine metabolism"

**Appendix 2.** **Additional compound targets for feature extraction (1).** Non-AMRT database metabolites with potential relevance to alkaptonuria (AKU) and/or tyrosine metabolism.

| Name | Formula | Mass | | CAS ID |
| --- | --- | --- | --- | --- |
| 1-(2-Carboxyphenylamino)-1'-deoxy-D-ribulose 5'-phosphate | C12H16NO9P | 349.056268 | | 5962-18-5 |
| 2,5-Dihydroxybenzaldehyde | C7H6O3 | 138.031694 | | 1194-98-5 |
| 2-Aminomuconic acid semialdehyde | C6H7NO3 | 141.042593 | |  |
| 2-Dehydro-3-deoxy-D-arabino-heptonate 7-phosphate (DAHP) | C7H13O10P | 288.024633 | | 2627-73-8 |
| 3,4-Dihydroxymandelaldehyde | C8H8O4 | 168.042259 | | 13023-73-9 |
| 3-Dehydroquinic acid | C7H10O6 | 190.047738 | | 10534-44-8 |
| 3-Hydroxytyrosol | C8H10O3 | 154.06299 | |  |
| 3-Hydroxytyrosol sulfate | C8H10O6S | 234.01981 | |  |
| 3-Methoxy-4-hydroxymandelate | C9H9O5 | 197.0449984 | |  |
| 3-Methoxy-4-hydroxyphenylglycolaldehyde | C9H10O4 | 182.057909 | | 17592-23-3 |
| 4-Hydroxyphenylacetaldehyde | C8H8O2 | 136.052429 | | 7339-87-9 |
| 4-Hydroxyphenylpyruvic acid | C9H8O4 | 180.042259 | | 156-39-8 |
| 5-O-(1-Carboxyvinyl)-3-phosphoshikimate | C10H13O10P | 324.024633 | | 74708-67-1 |
| 5-phosphoribosyl-1-diphosphate | C5H13O14P3 | 389.951815 | | 7540-64-9 |
| Acetic acid | C2H4O2 | 60.021129 | | 64-19-7 |
| Acetyl-CoA | C23H38N7O17P3S | 809.125773 | |  |
| Acetyl-L-tyrosine | C11H13NO4 | 223.084458 | | 537-55-3 |
| Adenosine triphosphate (ATP) | C10H16N5O13P3 | 506.995745 | | 987-65-5 |
| Aminomuconic acid | C6H7NO4 | 157.037508 | |  |
| Chorismic acid | C10H10O6 | 226.047738 | | 617-12-9 |
| Coenzyme A (CoA) | C21H36N7O16P3S | 767.115208 | | 85-61-0 |
| Dehydro-HGA | C8H6O4 | 166.02661 | |  |
| D-Erythrose 4-phosphate | C4H9O7P | 200.008589 | | 585-18-2 |
| D-Glyceraldehyde 3-phosphate | C3H7O6P | 169.998024 | | 591-57-1 |
| D-Pantetheine 4'-phosphate | C11H23N2O7PS | 358.096358 | | 2226-71-3 |
| Glutamyl-Tyrosine | C14H18N2O6 | 310.116486 | |  |
| Hipposudoric acid | C16H8O8 | 328.02192 | |  |
| Indole | C8H7N | 117.057849 | | 120-72-9 |
| Indoleglycerol phosphate | C11H14NO6P | 287.055874 | | 4220-97-7 |
| Metanephrine | C10H15NO3 | 197.105193 | | 5001-33-2 |
| N-(5-Phospho-D-ribosyl)anthranilate | C12H16NO9P | 349.056268 | | 27695-85-8 |
| NADH | C21H29N7O14P2 | 665.1247717 | |  |
| NADPH | C21H30N7O17P3 | 745.091102 | | 2646-71-1 |
| N-Methyltyramine | C9H13NO | 151.099714 | | 370-98-9 |
| Norhipposudoric acid | C15H8O6 | 284.03209 | |  |
| Oxaloacetate | C4H4O5 | 132.005873 | | 328-42-7 |
| Phenylpyruvic acid | C9H8O3 | 164.047344 | | 156-06-9 |
| Phloretic acid | C9H10O3 | 166.062994 | | 501-97-3 |
| Prephenic acid | C10H10O6 | 226.047738 | | 126-49-8 |
| Pyrophosphate | H4O7P2 | 177.943225 | | 2466-09-3 |
| Pyruvate | C3H4O3 | 88.016044 | | 127-17-3 |
| S-Acetylphosphopantetheine | C13H25N2O8PS | 400.106923 | |  |
| S-Adenosylmethionine | C15H23N6O5S | 399.145064 | | 29908-03-0 |
| Salicyluric acid | C9H9NO4 | 195.05316 | |  |
| Shikimate-3-phosphate | C7H11O8P | 254.019154 | | 63959-45-5 |
| Tyramine glucuronide | C14H19NO7 | 313.116152 | | 27972-85-6 |
| Tyramine-O-sulfate | C8H11NO4S | 217.040879 | | 30223-92-8 |
| Tyrosol | C8H10O2 | 138.06808 | |  |
| Tyrosol 4-sulfate | C8H10O5S | 218.024894 |  | |
