## Appendix 3 for "Studies in alkaptonuria reveal new roles beyond drug clearance for phase I and II biotransformations in tyrosine metabolism"

**Appendix 3.** **Additional compound targets for feature extraction (2).** Non-AMRT database metabolite products from predicted phase I and II biotransformations of homogentisic acid (HGA).

| Name | Formula | Mass | Phase |
| --- | --- | --- | --- |
| HGA 1,4-Dihydropyridines to Pyridines | C8H6O4 | 166.02661 | I |
| HGA 1,4-Dihydropyridines to Pyridines | C8H6O4 | 166.02661 | I |
| HGA Alcohols Dehydration | C8H6O3 | 150.03169 | I |
| HGA Alkene to Epoxide | C8H8O5 | 184.03717 | I |
| HGA Alkenes to Dihydrodiol | C8H10O6 | 202.04774 | I |
| HGA Decarboxylation | C7H8O2 | 124.05243 | I |
| HGA Deethylation | C6H4O4 | 140.01096 | I |
| HGA Demethylation | C7H6O4 | 154.02661 | I |
| HGA Demethylation and Hydroxylation | C7H6O5 | 170.02152 | I |
| HGA Demethylation and Methylene to Ketone | C7H4O5 | 168.00587 | I |
| HGA Demethylation and two Hydroxylations | C7H6O6 | 186.01644 | I |
| HGA Ethyl Ether to Acid | C6H2O5 | 153.99022 | I |
| HGA Ethyl to Carboxylic Acid | C7H4O6 | 184.00079 | I |
| HGA Hydration, Hydrolysis (Internal) | C8H10O5 | 186.05282 | I |
| HGA Hydroxylation and Desaturation | C8H6O5 | 182.02152 | I |
| HGA Hydroxymethylene Loss | C7H6O3 | 138.03169 | I |
| HGA Isopropyl to Acid | C6H2O6 | 169.98514 | I |
| HGA Ketone to Alcohol | C8H10O4 | 170.05791 | I |
| HGA Tert-Butyl to Acid | C5O6 | 155.96949 | I |
| HGA (O, N, S) Methylation | C9H10O4 | 182.05791 | II |
| HGA 2x Sulfate Conjugation | C8H8O10S2 | 327.95589 | II |
| HGA Acetylation | C10H10O5 | 210.05282 | II |
| HGA Cysteine Conjugation | C11H15NO6S | 289.06201 | II |
| HGA Cysteine Conjugation and Desaturation | C11H13NO6S | 287.04636 | II |
| HGA Cysteine Glycine Conjugation | C13H18N2O7S | 346.08347 | II |
| HGA Glucuronide Conjugation | C14H16O10 | 344.07435 | II |
| HGA Glutamine Conjugation | C13H16NO6 | 282.09776 | II |
| HGA Glycine Conjugation | C10H11NO5 | 225.06372 | II |
| HGA Hydroxylation + Glucuronide | C14H16O11 | 360.06926 | II |
| HGA Hydroxylation and Methylation | C9H10O5 | 198.05282 | II |
| HGA Hydroxylation and Sulfation | C8H8O8S | 263.99399 | II |
| HGA N-Acetylcysteine Conjugation | C13H17NO7S | 331.07257 | II |
| HGA N-Acetylcysteine Conjugation and Desaturation | C13H15NO7S | 329.05692 | II |
| HGA Sulfate Conjugation | C8H8O7S | 247.99907 | II |
| HGA Sulfate Conjugation | C8H8O7S | 247.99907 | II |
| HGA Taurine Conjugation | C10H13NO6S | 275.04636 | II |
