## Appendix 4 for "Studies in alkaptonuria reveal new roles beyond drug clearance for phase I and II biotransformations in tyrosine metabolism"

**Appendix 4.** **Non-targeted feature extraction parameters.**

Molecular feature extraction (MFE) parameters for ‘recursive non-targeted feature extraction’ were as follows: peak height >5000 counts, charge state range 1-2, minimum ion count = 1, binning and alignment tolerances - RT window = 0.3 min and mass window = 50 ppm, MFE score >70 in 60% of samples in at least one sample group. ‘Find by ion’ parameters (as part of the recursive feature extraction workflow) were as follows: mass score = 100, isotope abundance score = 60, isotope spacing score = 50, RT score = 100, expected data variation = 50 ppm (15%) and RT 0.3 min, integration - ‘Agile 2’ algorithm, extracted ion chromatogram (EIC) smoothed before integration, smoothing function - Gaussian, peaks filtered by height (absolute height >1000 counts), spectra to include - average scans >40% of peak height, exclude if >20% of saturation, centroiding - maximum spike width = 2, required valley = 0.7, ‘find by ion’ filters - score >60 in at least one sample group. Allowed ion species and sample groupings were the same as described for targeted data mining.
