## Appendix 5 for "Studies in alkaptonuria reveal new roles beyond drug clearance for phase I and II biotransformations in tyrosine metabolism"

**Appendix 5. Number of entities retained before and after quality control (QC).** QC filtering using pooled group sample data was performed in Mass Profiler Professional. **‘**Targeted’ and ‘non-targeted’ refer to feature extraction approaches. Entities from targeted feature extraction are AMRT-matched against a database generated from IROA Technology MS metabolite library of standards (466 compounds) or accurate mass-matched against an appended list of compounds from wider tyrosine metabolism (85 metabolites). Entities from non-targeted feature extraction are unidentified.

|  | *Hgd*^-/-^ vs *Hgd*^+/-^ | |  | |
| --- | --- | --- | --- | --- |
|  | Targeted | | Non-targeted | |
| QC filtering step | Positive | Negative | Positive | Negative |
| Manual curation of entities only | 323 | 324 | 662 | 552 |
| Present in 100% of replicate injections of at least one pooled group sample | 323 | 324 | 569 | 525 |
| CV <25% across replicate injections of each pooled group sample | 243 | 250 | 213 | 359 |
